# A Canine Model of Chronic Ischemic Heart Failure

**DOI:** 10.1101/2022.10.07.511342

**Authors:** Muhammad S. Khan, Douglas Smego, Yuki Ishidoya, Annie M. Hirahara, Emmanuel Offei, Sofia R. Castillo, Omar Gharbia, Joseph A. Palatinus, Lauren Krueger, TingTing Hong, Guillaume L. Hoareau, Ravi Ranjan, Craig Selzman, Robin Shaw, Derek J. Dosdall

## Abstract

**Background:** There is a paucity of novel therapeutics for chronic heart failure (HF). Preclinical large animal models of chronic HF are crucial to translating fundamental discoveries. Canine models of ischemic cardiomyopathy are frequently utilized but are severely limited by competing needs to both induce sufficient muscle injury which results in an appropriately severe degree of left ventricular dysfunction yet to also minimize early mortality.

**Methods:** Twenty-eight healthy adult dogs (30±4 kg, 15/28 male) underwent thoracotomy procedures followed by one of three types of left anterior descending (LAD) coronary artery ligations: Simple LAD (proximal and distal LAD ligation), LAD and Lateral (proximal and distal LAD, distal first diagonal, and proximal first obtuse marginal), and Total LAD Devascularization (TLD including the proximal and distal LAD, proximal aspects of the first, second, and third diagonals, and proximal aspects of first and second epicardial right ventricular branches of the LAD). Invasive and non-invasive hemodynamics were followed until attainment of chronic severe HF, which was achieved when left ventricular ejection fraction (LVEF) was <40% and N-terminal prohormone B-type Natriuretic peptide (NT-proBNP) >900 pmol/L, both for two consecutive measurements.

**Results:** Overall post-ligation early mortality (within 48 h of coronary ligation) was 18% (n=5/28), and the overall survival rate at 5 weeks post-ligation was 68% (n=19/28). At post-ligation week 1, LVEF was acutely reduced in all animals, from 54±4 to 38±1%, p<0.001. Beyond week 1, systolic function recovered over 3-4 weeks and then slowly declined as the left ventricle dilated and both systolic and diastolic function worsened. The magnitude of the initial decline in LVEF predicted ultimate HF development. Dogs with Simple LAD and LAD plus Lateral had excellent and limited survival at 48 hours (100% and 47%, respectively), yet the second group more likely developed chronic HF at 9 – 14 weeks (50% versus 57%, respectively). Dogs with TLD had both improved survival at 48 hours (90%) and excellent development of chronic HF (100%).

**Conclusions:** By focusing on the LAD only and limiting collateral flow to it by ligating all side branches, optimal survival and attainment of chronic ischemic HF can be achieved in the canine heart.

## INTRODUCTION

Preclinical large animal HF models play a critical and expanding role in translating basic science findings to development and approval of novel cardiovascular therapeutics.^1^ While small animal models are generally cheaper, allow for larger sample sizes, and offer a greater variety of transgenic models,^2^ they have significant limitations in HF with reduced ejection fraction (HFrEF) research including small organ and body size, faster heart rates, different action potential characteristics, and different myocardial metabolism.^3^ In vivo studies using large mammalian hearts are advantageous because they are more physiologically and clinically relevant,^4^ and permit testing of clinically relevant interventions.^5^

Canines and swine are the most frequently used large animal species for chronic heart failure studies, and the hearts of both animals exhibit human characteristics.^6^ Coronary circulation of pigs has no significant collateral circulation,^7^ while the canine model has epicardial collateral circulation.^8^ The collateral circulation of dogs more closely replicates patient hearts with ischemic heart disease (which promotes collateral growth), especially in older individuals.^8^ The social nature of dogs often eliminates the need for sedation for minimally-invasive procedures such as transport, echocardiography, electrocardiogram monitoring, and phlebotomy. A limitation of dog use in chronic infarction models is that collateral circulation can make it difficult to induce consistent myocardial infarctions of sufficient size that heart failure develops. If too much coronary circulation is impaired, then post-ligation mortality of canines is unacceptably high, between 30 and 60%^9–13^, which similar to that of the porcine infarction models.^8^

An optimized approach for developing chronic ischemic heart failure in a canine model will minimize early mortality and result in a repeatable infarct size that leads to chronic dilated cardiomyopathy. In this work, we compare various coronary ligation procedures and test the hypothesis that elimination of the epicardial collateral circulation will decrease the acute loss rate and improve the consistency of heart failure progression in the dog model. We present a new approach of Total Left anterior descending Devascularization (TLD), which addresses these concerns and provides a repeatable, consistent model of ischemic heart failure in canines. This model will be a valuable tool for pre-clinical testing of therapeutics and medical devices and basic research in HFpEF and ischemic HF,

## METHODS

### Animal and Surgical Preparation

All studies followed the guidelines from the National Institutes of Health *Guide for the Care and Use of Laboratory Animals*^14^ and standards of United States regulatory agencies. Protocols were approved by the Institutional Animal Care and Use Committee (IACUC) of The University of Utah. The dogs with baseline (BL) left ventricular ejection fraction (LVEF) >40% and N-terminal prohormone B-type Natriuretic peptide (NT-proBNP) < 900 pmol/L were included in the study. Studies were performed on 28 healthy adult dogs (female, n=14) weighing 30±4 kg (25.4 – 39.6). Dogs were fasted for 12 – 18 h, and a fentanyl patch (50 – 100 mcg/h, dermal) was placed 3 – 12 h before the infarct surgical procedure. Dogs were anesthetized with an intravenous (IV) injection of 1.5 – 2 mL fentanyl followed by Propofol (5 – 8 mg/kg, IV). The animal was then intubated, and anesthesia was maintained with a mixture of 1.0 – 3.0% isoflurane in oxygen through a ventilator. Heart rate (HR), core body temperature, oxygen saturation, and electrocardiogram were monitored. Expired pCO2 was monitored and corrected when necessary by adjusting tidal volume and respiratory rate during the procedures. Once the animal was hemodynamically stable under general anesthesia, fentanyl was administered at 2-10 mcg/(kg*hr), and Cefazolin (20 mg/kg, IV) was also given every 90 – 120 minutes. Intraoperative hypotension (mean arterial pressure < 60 mmHg) was treated with a bolus of Lactated Ringer’s solution (75 – 200 mL, IV) followed by an infusion of dopamine (2-10 mcg/(kg*min). Atropine (0.01 – 0.05 mg/kg, IV bolus) was administered for sinus bradycardia and low blood pressure (BP) (mean < 60 mmHg) to restore normal cardiac function and heart rate. A femoral catheter was used for invasive BP monitoring, and the other femoral artery was cannulated for PV loop catheter access.

### Thoracotomy Procedure

Before the thoracotomy procedure, bupivacaine (1 – 2 mg/kg, SC) was given at the thoracotomy site, and a nerve block was induced by injection into the intercostal spaces. The heart was exposed by incising the skin, followed by underlying layers parallel to the rib of the 5^th^ intercostal space. As shown in **Fig. 1**, the lateral anterior descending (LAD) artery and collateral vessels were isolated and ligated in three groups. Group 1 (**Simple LAD,** n=4): ligations as the proximal and distal LAD, with a ligation above the first diagonal of the LAD and a second ligation at the junction of the distal LAD and the collateral connection to the posterior descending artery near the apex of the LV. Group 2 (**LAD and Lateral,** n=15): ligations at the proximal and distal LAD, distal end of the first diagonal where there may be a collateral connection between the LAD perfusion bed and the first obtuse marginal perfusion bed, and proximal first obtuse marginal artery. Group 3 (**Total LAD Devascularization, TLD,** n=9): ligations at the proximal and distal LAD, proximal aspects of visible diagonal arteries off the LAD, and also proximal aspects of visible epicardial right ventricular branches of the LAD below the first diagonal artery. Once completed, a chest tube was inserted before closing the thoracotomy site. The chest tube was then used to evacuate the air and any fluid from the thorax. The tube was finally removed before the dog was returned to the recovery kennel. After the surgical procedure, bupivacaine (1 mg/kg, IM/SQ) or Nocita (5.3 mg/kg maximum dose, IM/SQ) was again administered at the incision site.

**Figure 1.**
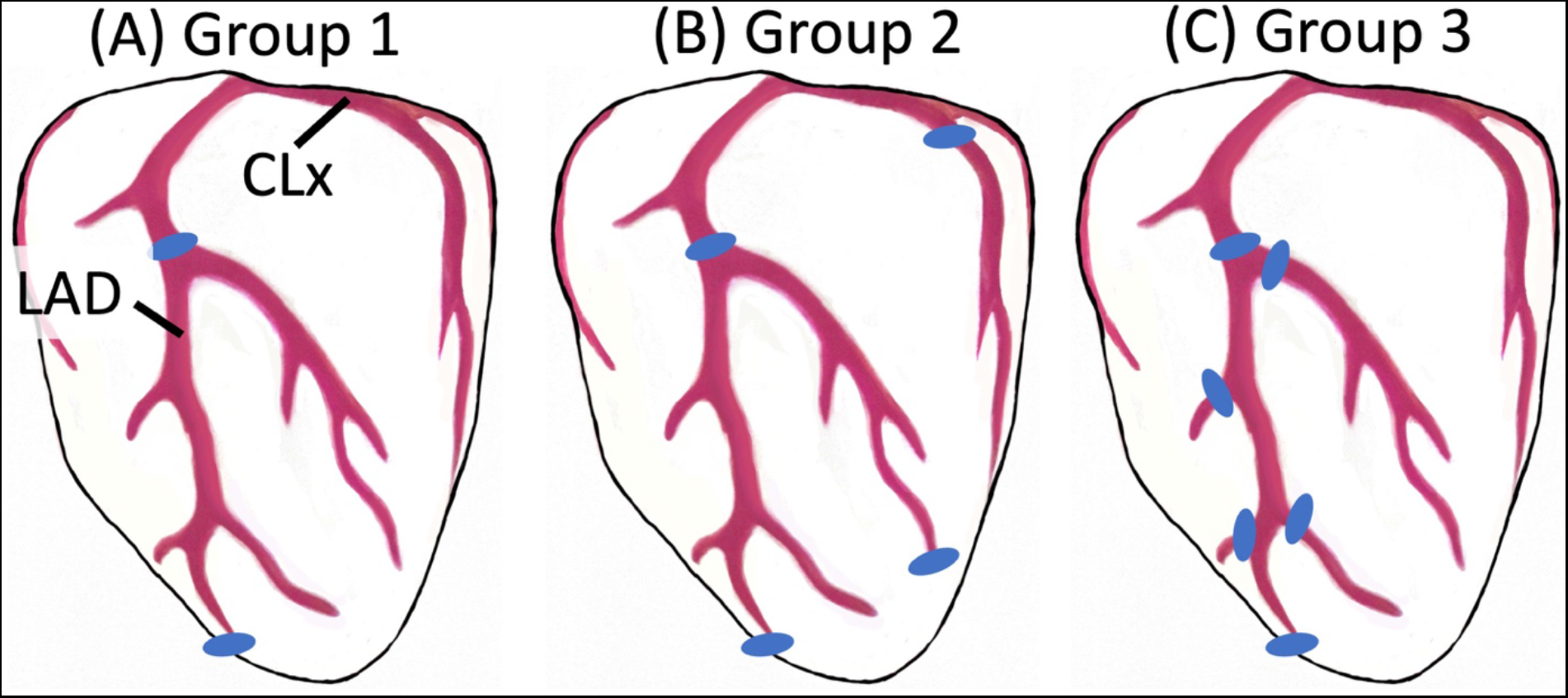
Coronary LAD artery ligation positions of the three groups. Ligation locations shown in blue. Group 1: Simple LAD (proximal and distal LAD ligation). Group 2: LAD and Lateral (proximal and distal LAD, distal first diagonal, and proximal first obtuse marginal). Group 3: Total LAD Devascularization (TLD including the proximal and distal LAD, proximal aspects of visible diagonals, and proximal aspects of epicardial right ventricular branches of the LAD). LAD: left anterior descending coronary artery. LCx: left circumflex artery.

### Antiarrhythmic Drugs and Protocols

In group 1, none of the dogs received any anti-arrhythmic drugs before the baseline procedure. For group 2, four dogs were given amiodarone (5 – 10 mg/kg, PO, BID) for a week before the baseline procedure, and continued for up to 1-week post-ligation period if the animal experienced a substantial number of premature ventricular contractions (PVCs) and non-sustained VT. For group 3, all animals received mexiletine (3 – 8 mg/kg PO TID) beginning the evening before the baseline procedure, and continued for 3 days post-ligation. A bolus of lidocaine (0.8 – 2.0 mg/kg, IV) was used during the procedure and / or post-operative care to control acute arrhythmias if they occurred. Once the lidocaine bolus effectively reduced arrhythmias, a continuous infusion of lidocaine (25 – 100 mcg/(kg*min), IV) was then used, if required, up to a cumulative total of 8 mg/kg, to reduce runs of VT. During recovery from anesthesia, hydromorphone (0.05 – 0.3 mg/kg, IM, SQ) was also administered. Finally, a magnesium bolus (0.1 mmol/kg, IV) was administered, if needed, during postoperative 2-3 days to control intractable arrhythmias.

### Let Ventricular End-Diastolic Pressure

For left ventricular end-diastolic pressure (LVEDP) measures, the present study used 7F combined pressure conductance catheter (CD Leycom, Zoetermeer, Netherlands) introduced retrogradely from the femoral artery into the LV via the aortic valve before performing LAD coronary ligation. The catheter was connected to a Cardiac Function Laboratory (CD Leycom) to acquire pressure data. All recordings were made during sinus beats with and with mechanical ventilation suspended. Data were exported and analyzed in a custom-built MATLAB program to evaluate LVEDP. LVEDP measurement was performed at the final study.

### Echocardiography Recordings and Analysis

LV function was assessed by echocardiography (Acusion SC2000, Siemens, Munich, Germany) in awake dogs before the baseline coronary ligation procedure and throughout HF development. Data were acquired weekly (BL to 4 weeks post-ligation) and biweekly (>4 weeks post-ligation) to evaluate both systolic and diastolic dysfunction over time. Data were gathered with apical views (4C, 3C, and 2C), parasternal long-axis view, and short-axis views: base (mitral valve level), mid (papillary muscle level), and apex. E and e’ peaks were recorded using pulse wave (PW) with tissue Doppler imagining (TDI) and PW alone, respectively, from a 4-chamber apical view. The EPSS was obtained by placing the M-mode tracer over the distal tip of the mitral valve’s anterior leaflet. The image displayed the movement of structures over several cardiac cycles along a specific plane. Data spanning at least three consecutive heartbeats were acquired and stored as digital images.

### Chronic Ischemic HF Criteria

Dogs were classified as achieving chronic ischemic HF if LVEF <40% and NT-proBNP >900 pmol/L for at least two consecutive measurements. Data were studied and characterized at different time-points as baseline procedure before thoracotomy (BL), chronic HF progression (week 1 to 16), and the final end-point (chronic ischemic HF developed during the HF progression timeline).

### Statistical Analysis

Data are expressed as mean ± standard deviation (SD). A paired t-test was used to compare the baseline data with post-ligation data (week 1, week 2, week 3, week 4, week 6, and final end-point) recorded serially at each time point in the same group. An unpaired t-test compared the baseline data with post-ligation data recorded among three groups (group 1, group 2, and group 3). A p-value below 0.05 and 0.01 were considered statistically significant and highly significant, respectively.

## RESULTS

A total of twenty-nine dogs subjected to LAD coronary ligation surgical procedures. One dog was lost due to anesthetic complications during the baseline study before the thoracotomy procedure and was excluded from the study. Among the 28 dogs studied (female, n=14), five dogs experienced sudden cardiac death in their recovery kennel within 48 h after the coronary ligation procedure, representing an early mortality rate of 18%. Another four dogs died in the first five weeks of HF progression of which three were euthanized: one dog experienced acute bleeding on post-ligation day 8; the second developed a pneumothorax on post-ligation day 4, and the third had intractable sustained ventricular tachycardia on post-ligation day 5. The fourth dog experienced sudden death in the kennel at week 5 post-ligation. The overall post-ligation 5-week survival rate was 68% (19/28), divided between 71% for females (n=10/14) and 64% for males (n=9/14).

The twenty-eight dogs were further categorized into three groups based on their different LAD coronary ligation surgical procedures, as shown in **Fig. 2** and **Table 1**. For group 1 and group 2, the post-ligation 5-week survival rate was 100% and 50%, respectively. However, in each group, the success rate of developing chronic ischemic HF was only 50%. The other 50% of survived dogs developed initial ischemic insult at week 1, as seen in **Fig. S1**, but failed to achieve the criteria of chronic ischemic HF. With the TLD group, group 3 had a 100% success rate of developing chronic ischemic HF (n=7/7) with an early mortality rate of 10% (**Table 1**).

**Table 1.**
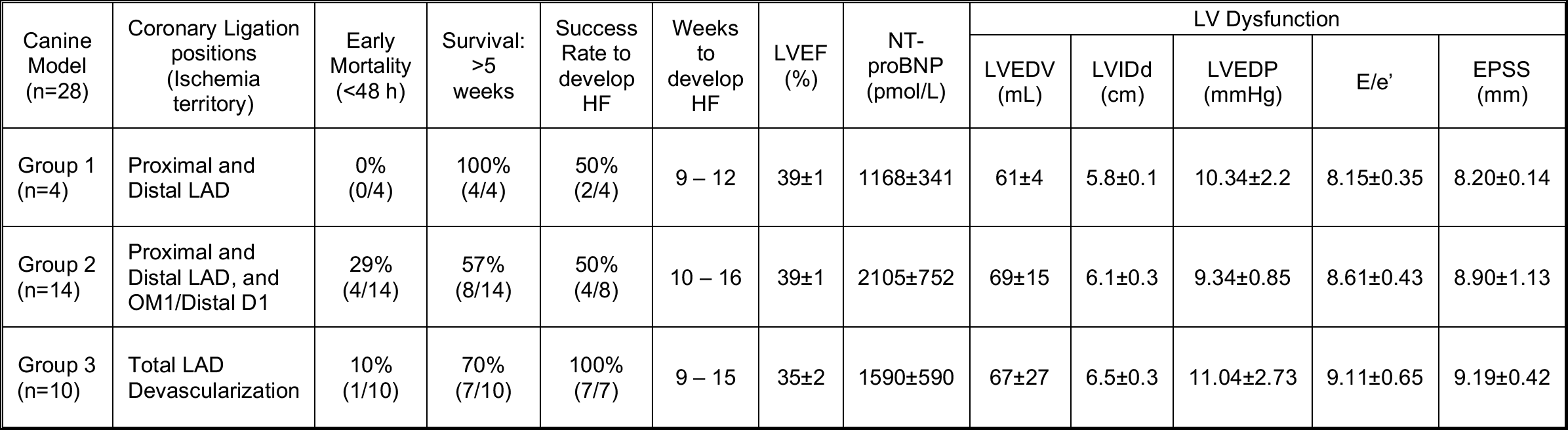
Survival rates and cardiac performance of three groups of dogs in developing chronic ischemic HF. LVEF: left ventricular ejection fraction. NT-proBNP: N-terminal prohormone B-type Natriuretic peptide. LVEDV: left ventricular end-diastolic volume. LVEPD; left ventricular diastolic pressure. LVIDd: left ventricle internal diastolic diameter. E: Early mitral inflow velocity. e’: Mitral annular early diastolic velocity. EPSS: End-point septal separation.

**Figure 2.**
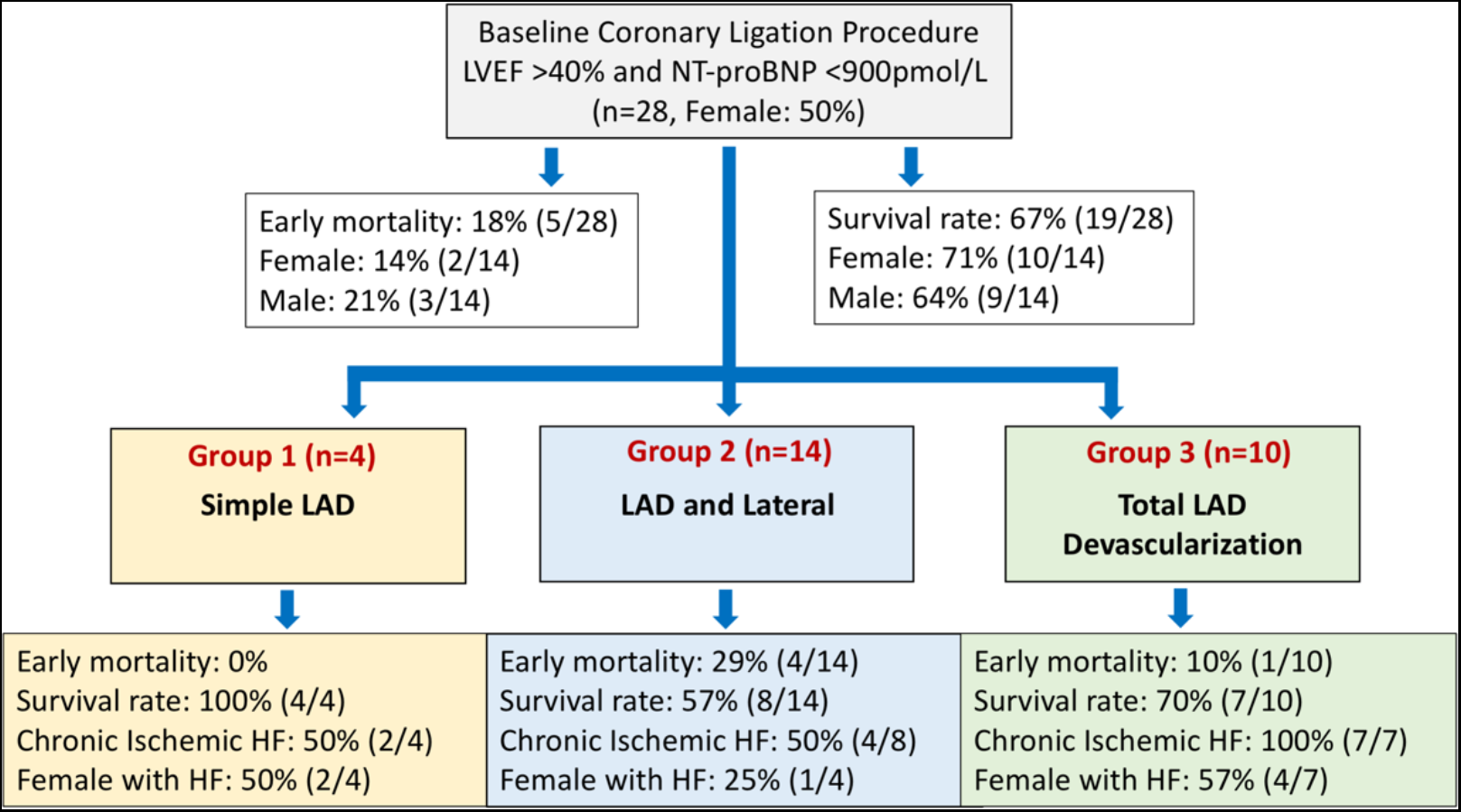
Flow chart description of healthy adult dogs enrolled in the study and the outcome of each group. Early mortality: <48 h post thoracotomy. Survival rate: Post-ligation >5 weeks. Chronic ischemic HF: LVEF <40% and NT-proBNP >900 pmol/L. LVEF: Left ventricular ejection fraction. NT-proBNP: N-terminal prohormone B-type Natriuretic peptide.

### LVEF and NT-proBNP

As seen in **Fig. 3a**, at post-ligation week 1, in group 1, group 2 and group 3, the LVEF decreased from 58±5% at baseline to 48±1% (ΔLVEF: ~23%), 54±2% to 42±2% (ΔLVEF: ~22%, p<0.01) and 50±3% to 40±3% (ΔLVEF: ~21%, p<0.01), respectively. Beyond week 1, the systolic function partially recovered over 3-4 weeks, as observed in **Fig. 3a**, followed by a slowed decline in LVEF. The decline in LVEF for Groups 2 and 3 were more dramatic and were statistically significant.

**Figure 3.**
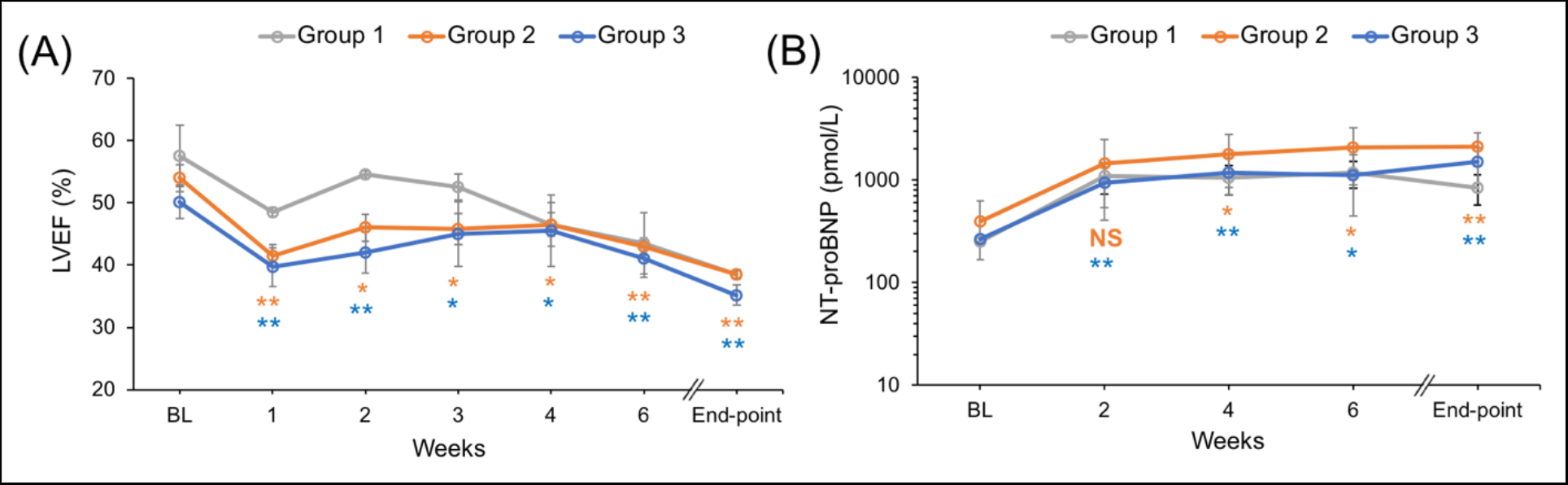
LVEF and serum NT-proBNP over time (Group 1: n=2, Group 2: n=4 and Group 3: n=7). (A) LVEF data acquired in 4-chamber apical view. (B) Plasma NT-proBNP level. ‘BL’ shows the data recorded before the baseline thoracotomy procedure in healthy dogs with LVEF >40%. ‘End-point’ represents final data time when dogs were determined to be in the chronic ischemic HF with LVEF <40% and NT-proBNP >900pmol/L. For groups 2 and 3, each time-point data was statistically compared with BL measures. For group 1, the presented data were averaged from the two dogs that developed chronic ischemic HF, and thus cannot be compared statistically. *p<0.05. **p<0.01. NS: not-significant. LVEF: left ventricular ejection fraction. NT-proBNP: N-terminal prohormone B-type Natriuretic peptide.

Plasma NT-proBNP was investigated along with LVEF to assess chronic HF progression. In 28 dogs, the NT-proBNP baseline level before coronary ligation was 322±155 pmol/L. As seen in **Fig. 3b**, plasma NT-proBNP was elevated in all groups at week 2 (1092±365, 1443±1036, 938±397 pmol/L), week 4 (1042±334, 1795±953, 1168±461 pmol/L) and at week 6 (1168±341, 2054±1159, 1113±664 pmol/L). Compared with pre-ligation plasma NT-proBNP level, the change at the final end-point when LVEF <40%, the NT-proBNP was significant in group 2 (2105±752 with p<0.01) and group 3 (1590±590 pmol/L with p<0.01). Finally, as observed in **Table 1** and **Fig. 2**, all groups that achieved the LVEF <40% at the final end-point had a plasma NT-proBNP >900 pmol/L.

### Left Ventricular Chamber Size and Diastolic Function

From pre-ligation to post-ligation as early as week 1, left ventricular internal diameter (LVIDd) increased in group 3 (5.4±0.5 vs. 5.5±0.5 cm, p<0.05). For group 2, a significant increase in LVIDd was observed at week 6 (5.9±0.3 cm, p<0.05) (**Fig. 3a**). At the time of attainment of chronic ischemic HF in group 2, the LVIDd had increased to 6.1±0.3 cm, p<0.05 whereas with the TLD (group 3), the LVIDd increased to 6.5±0.3 cm, p<0.01. Overall, the LVIDd percentage increase from the baseline to the final end-point in group 1 (7%) and group 2 (13%), were lower relative to that of group 3 (18%).

Diastolic function was evaluated by echocardiogram. Mitral E/e’ ratio (E: Early mitral inflow velocity, e’: Mitral annular early diastolic velocity) and end-point septal separation (EPSS) were measured to assess diastolic dysfunction in group 2 and group 3, as shown in **Fig. 4b** and **Fig. 4c**, respectively. For group 1, E/e’ and EPSS data were only recorded during the final endpoint. For group 3, compared to baseline measures (4.71±0.66 mm and 7.01±0.52), both EPSS and E/e’, respectively, increased significantly at week 1 (6.97±1.45 mm, p<0.01 and 8.33±0.85, p<0.01), week 2 (6.71±1.55 mm and 8.35±0.98, p<0.01), week 3 (7.44±0.94 mm and 8.51±0.90, p<0.01), week 4 (7.96±0.94 mm and 8.47±0.56), week 6 (8.60±0.58 mm, p<0.01 and 8.56±0.73, p<0.01) and at the final end-point (9.19±0.42 mm, p<0.01 and 9.11±0.65, p<0.01). For group 2, EPSS and E/e’ did not experience significant changes from week 1 to week 4, but did significantly increase at week 6 (8.02±0.36 mm, p<0.01 and 7.80±0.19, p<0.05) and at the final end-point (8.90±1.13 mm, p<0.01 and 8.61±0.43, p<0.05).

**Figure 4.**
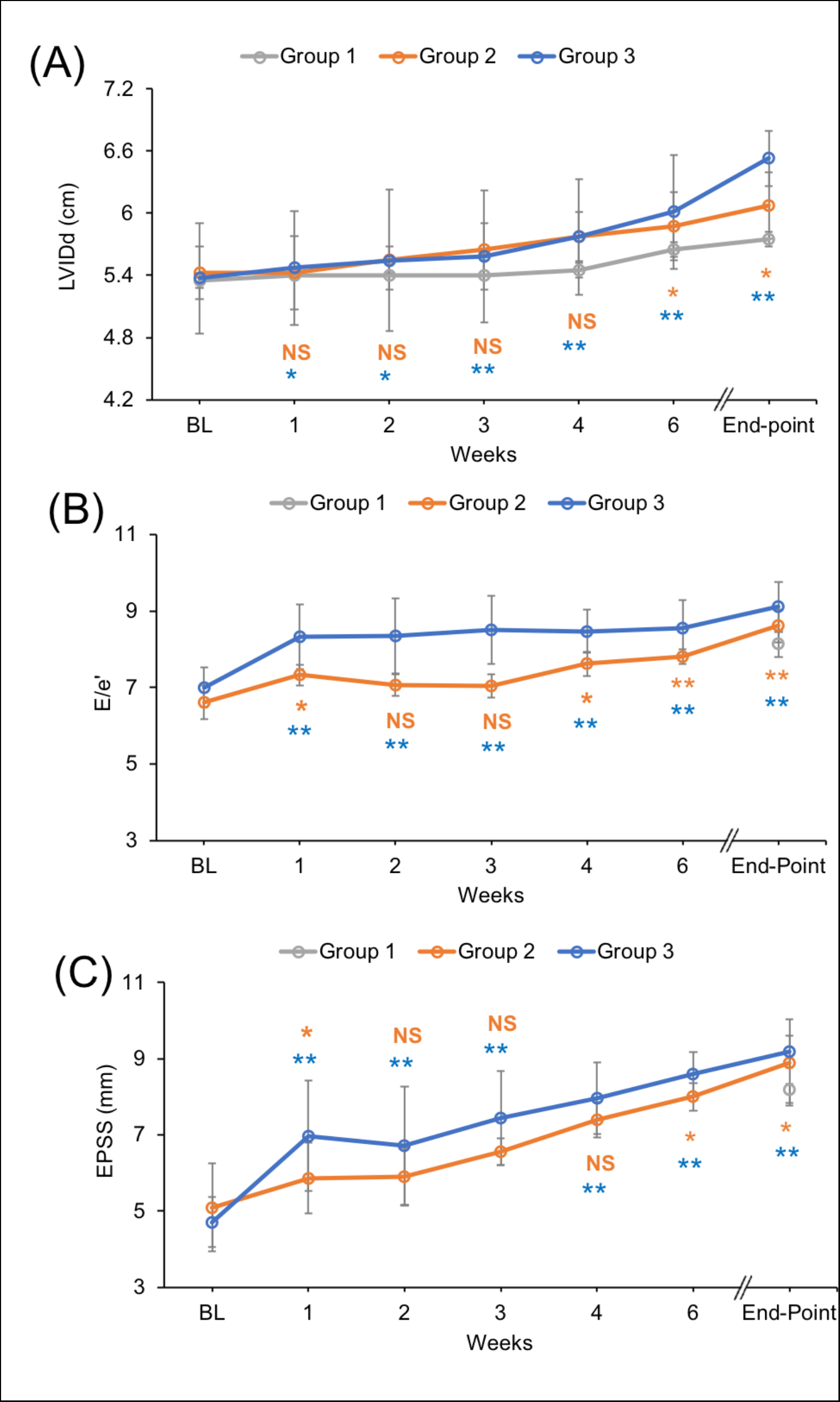
Left ventricular dysfunction during chronic ischemic HF progression in three groups. (A) LVIDd data acquired in 4-chamber apical view (Group 1: n=2, Group 2: n=4 and Group 3: n=7). (B) E/e’ data acquired and quantified through PW (TDI) images from a 4-chamber apical view. (C) EPSS data were recorded in m-mode from a right parasternal long axis view. For Group 1, in (B) and (C) the data points were only available at the final end-point. ‘BL’ shows data recorded before the baseline thoracotomy procedure, and ‘end-point’ represents final data time when dogs met criteria for chronic ischemic HF. For each time-point, data were statistically compared with BL measures except for the Group 1. *p<0.05. **p<0.01. NS: not-significant. LVIDd: left ventricle internal diastolic diameter. E: Early mitral inflow velocity. e’: Mitral annular early diastolic velocity. EPSS: End-point septal separation.

LVEDP was recorded prior to thoracotomy and at the final end-point surgical procedures in anesthetized dogs. As seen in **Fig. 5**, LVEDP increased in all groups at the final endpoint. Group 2 and group 3 exhibited a significant change in LVEPD from the baseline to the final endpoint: 5.22±0.22 to 9.34±0.85 mmHg, p<0.01, and 6.29±1.18 to 11.04±2.73 mmHg, p<0.01.

**Figure 5.**
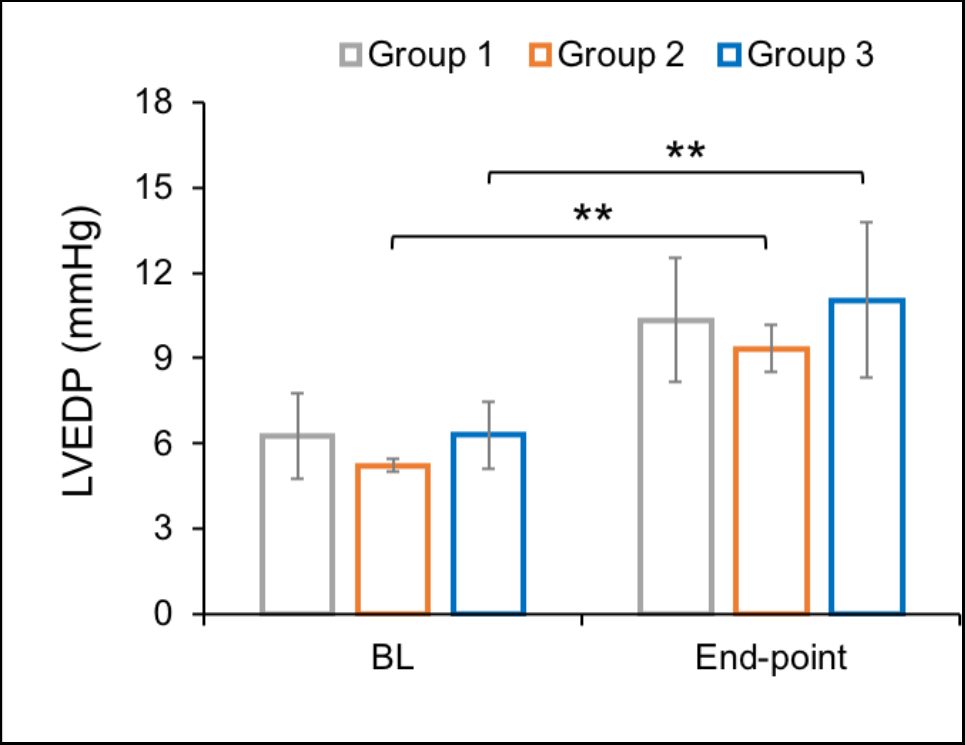
Left ventricle end-diastolic pressure (LVEDP) acquired from pressure-volume loops in three groups at baseline and at final end-point of chronic ischemic HF. **p<0.01.

## DISCUSSION

Often, as outlined in **Table 2**, a canine model of chronic ischemic HF was typically produced through single or multiple coronary microembolizations. The early mortality rate of ischemia by microembolization varies between 30% and 60%.^9–11,13,15^. Ameroid constrictors, which is another method to achieve ischemic cardiomyopathy, consists of placement of constrictors on proximal portions of the LAD and left circumflex (LCx) coronary arteries. The ameroid approach results in an early mortality of about 45%.^12^ In our chronic ischemic HF canine model, we reported an overall early mortality rate of 18% (n=5/27), and 30% (n=8/27) by post-ligation week 5. For group 1 (LAD) and group 2 (LAD plus lateral), the success rate of developing a robust chronic ischemic HF was 50%. In both techniques, the collateral flow was not restricted sufficiently and thus ischemic areas recovered with collateral flow whereas acutely infarcted areas lead to sudden arrhythmic death in group 2. For group 3, as shown in **Fig. 1**, TLD technique involved the maximum restriction of collateral blood flow across LAD perfusion bed and thus produced chronic ischemia with a 100% success rate of developing chronic HF.

**Table 2.** Previously reported chronic ischemic HF canine models. LVEF: left ventricular ejection fraction. NT-proBNP: N-terminal prohormone B-type Natriuretic peptide. LVEPD; left ventricular diastolic pressure, LVEDV: left ventricular end-diastolic volume. LVESV: left ventricular end-systolic volume. ‘Δ’ represents the change from the baseline to the final end-point once the model reached chronic ischemic HF condition. PNE: Plasma norepinephrine. LCx: left circumflex artery.

| Canine Model | Procedure Type to induce MI | Criteria to evaluate HF | Mortality rate | Weeks to develop HF | Reported parameters in developed chronic ischemic HF models |  |  |  | REF. |
| --- | --- | --- | --- | --- | --- | --- | --- | --- | --- |
|  |  |  |  |  | EF (%) | LVEDP (mmHg) | LV Infarct (%) | Others |  |
| n=20 | I/R – proximal and distal LAD and major diagonal branches | LVEF<40% | NA | 4 – 5 | <40 | 15±3.5 | 16.3±2.6 | NT-proBNP: 4081±1123pmol/L | 19 |
| n=20 | Single coronary microembolization into LCx | LVEF<35% | 55% | 44 | 23 | 14.9±2.5 | 23.1±2.6 | NA | 9 |
| n=17 | LAD coronary artery occlusion | LVEF<35% | NA | 4 – 5 | 31.7±4.6 | NA | 11±0.4 | PCWP: 18±1.2mmHg | 20 |
| n=22 | Ameroid constrictors placement on proximal LAD, LCx and their major branches | NA | 45% | 6 | NA | NA | NA | ΔLVEDA: 36% ↑, ΔLVESA: 86% ↑, ΔLV wall thickening: 57 ↓ | 12 |
| n=7 | Proximal and distal LAD and major diagonal branches | LVEF<35% | 0% | 8 – 10 | 28.7±5.4 | NA | NA | ΔLVEDV: 14% ↑, ΔLVESV: 25% ↑ | 21 |
| n=21 | Proximal LAD | LVEF>45% | 0% | 3 | >45 | ≥18 | NA | NA | 22 |
| n=7 | I/R – proximal and distal LAD, LCx and D1 LAD | NA | NA | 4 | 48±6 | 16.8±6.4 | 13.3±2.5 | NT-proBNP: 1778±195pmol/L | 23 |
| n=20 | Sequential coronary microembolization into LCx | LVEF<35% | 30% | 13 | 21±1 | 22±3 | 21±3 (anterior)<br>17±2 (posterior) | ΔLVEDV: 40% ↑, PNE: 791±131 pg/mL | 11 |
| n=32 | LAD coronary artery balloon occlusion | NA | 25% | 6 | NA | NA | 37.1±8.2 | NA | 15 |
| n=12 | Sequential coronary microembolization into LCx | LVEF<35% | NA | 17 – 18 | 23±2 | 22±3 | 22±3 | ΔLVEDV: 35% ↑, ΔLVESV: 144% ↑ | 24 |
| n=7 | Sequential coronary microembolization into LCx | LVEF:30 – 40% | NA | 18 | 26±1% | 20±5 | NA | ΔLVEDV: 28% ↑, PNE: 569±44 pg/mL | 25 |
| n=15 | Sequential coronary microembolization into LCx | LVEF<40% | 60% | 8 | 35±3% | NA | NA | ΔLVEDV: 22% ↑, PNE: 715±276 pg/mL | 13 |
| n=8 | Sequential coronary microembolization into LCx | LVEF≤35% | NA | 16 | 33% | NA | NA | ΔLVEDV: 11% ↑, ΔLVESV: 17% ↑ | 26 |

In all groups, in the first month post-ligation, LVEF rapidly declined to be followed by partial recovery as surviving animals transitioned from acute ischemic HF to a more chronic ischemic HF. After the first month, the hearts began to dilate and LVEF slowly declined again, with biomarkers revealing increased cardiac stretch. The dogs experienced a partial recovery of LV function after acute coronary occlusion, as has been previously reported.^16^ This restoration of LV function may be due to subsequent recovery of stunned myocardium with increased collateral flow, i.e. transient rather than permanent initial injury to cardiomyocytes.^17^ Beyond week 1, systolic function recovered over 3-4 weeks and then slowly declined as the LV dilated and both systolic and diastolic function worsened. The slower secondary decline is consistent with cardiac remodeling and the slow development of chronic HF. Ischemic but still viable myocardium is known to be highly arrhythmogenic.^18^ Finally, the criteria for severe chronic HF was achieved 2-4 months post ligation. It is interesting that TLD improved mortality, yet this form of coronary ligation resulted in larger loss of LV function at week 1 (**Fig. 3**) and increased attainment of chronic HF (**Table 1**). By causing a complete infarction of the LAD perfusion bed, along with improved survival we also achieve a higher rate of chronic HF attainment. TLD appears to be an optimized approach to, in canine animal models, achieving chronic ischemic cardiomyopathy.

In summary, the data obtained in this study indicate that a complete LAD coronary ligation together with ligation of collateral circulation branches, can create a robust canine model of chronic ischemic HF that is optimal for both animal survival and attainment of dilated cardiomyopathy. Based-on reported scientific evidence in our study, it indicates that the TLD approach in canines is highly reproducible with a 100% success in developing a robust pre-clinical ischemic HF.

## Conflict of Interest

RR is a consultant for Abbott, Medtronic, and Biosense Webster. All other authors have no relevant financial or non-financial interests to disclose.

## Funding

The research has been supported through awards: R01HL128752 (DJD), R21HL156039 (DJD), R21AG074593 (RMS and TH), R01HL152691 (RMS) and research grants from the Nora Eccles Treadwell Foundation (DJD, RMS). MSK is supported through NHLBI training grant 5T32HL007576-035.

---

**Figure S1.**
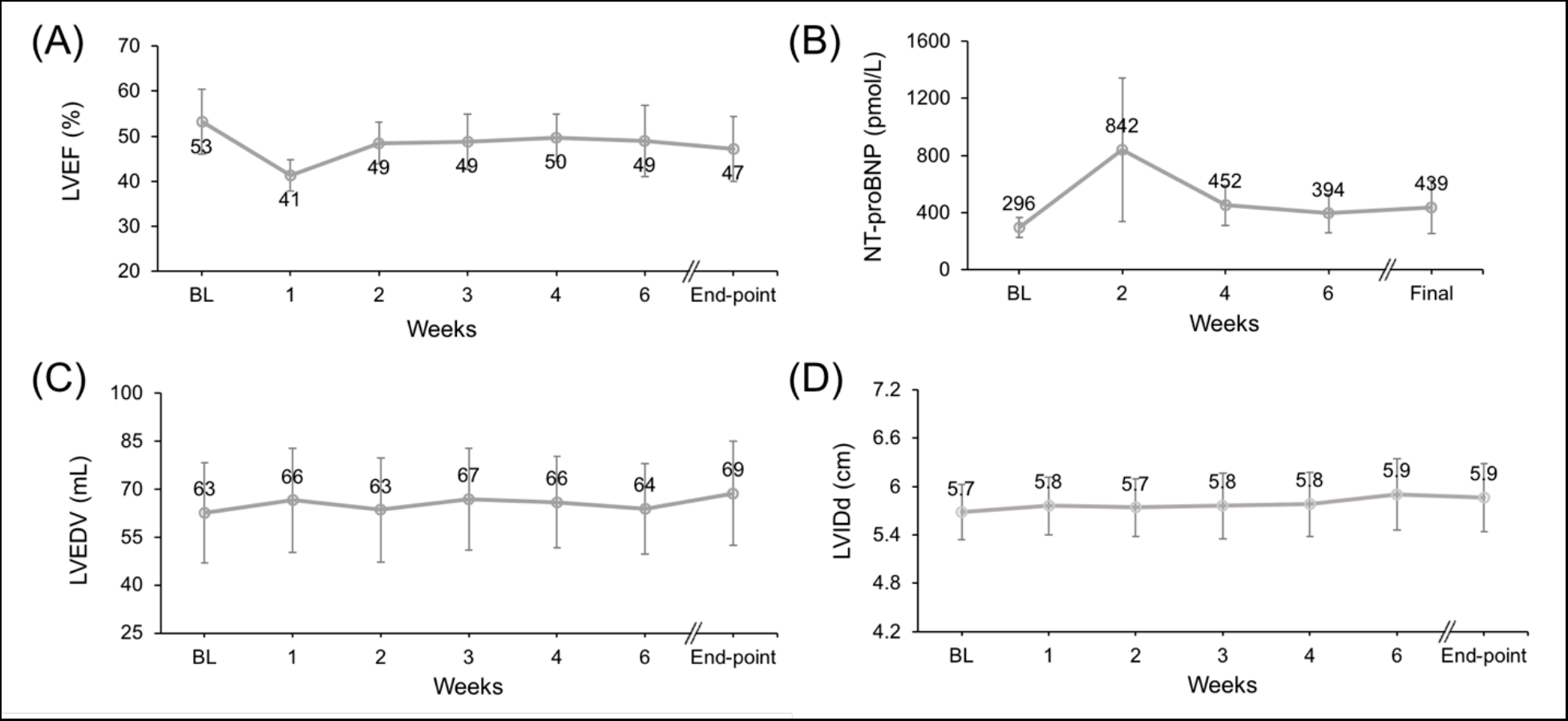
LV hemodynamic changes over time in a group of healthy adult canine models (n=5, female: 3/5) that did not reach criteria for chronic HF. Among five dogs, the two were from group 1 (female: 1/2) and the three were from group 2 (female: 2/3).

## References

(1) Lindsey, M. L.; Bolli, R.; Canty, J. M.; Du, X. J.; Frangogiannis, N. G.; Frantz, S.; Gourdie, R. G.; Holmes, J. W.; Jones, S. P.; Kloner, R. A.; et al. Guidelines for Experimental Models of Myocardial Ischemia and Infarction. American Journal of Physiology - Heart and Circulatory Physiology 2018, 314 (4), H812–H838.

(2) Riehle, C.; Bauersachs, J. Small Animal Models of Heart Failure. Cardiovascular Research 2019, 115 (13), 1838–1849.

(3) Ginis, I.; Luo, Y.; Miura, T.; Thies, S.; Brandenberger, R.; Gerecht-Nir, S.; Amit, M.; Hoke, A.; Carpenter, M. K.; Itskovitz-Eldor, J.; et al. Differences between Human and Mouse Embryonic Stem Cells. Developmental biology 2004, 269 (2), 360–380.

(4) Dixon, J. A.; Spinale, F. G. Large Animal Models of Heart Failure; A Critical Link in the Translation of Basic Science to Clinical Practice. Circulation: Heart Failure 2009, 2 (3), 262–271.

(5) Hearse, D. J.; Sutherland, F. J. Experimental Models for the Study of Cardiovascular Function and Disease. Pharmacological research 2000, 41 (6), 597–603.

(6) Spannbauer, A.; Traxler, D.; Zlabinger, K.; Gugerell, A.; Winkler, J.; Mester-Tonczar, J.; Lukovic, D.; Müller, C.; Riesenhuber, M.; Pavo, N.; et al. Large Animal Models of Heart Failure With Reduced Ejection Fraction (HFrEF). Frontiers in Cardiovascular Medicine 2019, 6 (17), 1–8.

(7) Recchia, F. A.; Lionetti, V. Animal Models of Dilated Cardiomyopathy for Translational Research. Veterinary research communications 2007, 31 *Suppl 1* (SUPPL. 1), 35–41.

(8) White, F. C.; Roth, D. M.; Bloor, C. M. The Pig as a Model for Myocardial Ischemia and Exercise. Laboratory animal science 1986, 36 (4), 351–356.

(9) Franciosa, J. A.; Heckel, R.; Limas, C.; Cohn, J. N. Progressive Myocardial Dysfunction Associated with Increased Vascular Resistance. American Journal of Physiology - Heart and Circulatory Physiology 1980, 239 (4), H477–H482.

(10) Smiseth, O. A.; Lindal, S.; Mjøs, O. D.; Vik-Mo, H.; Jørgensen, L. Progression of Myocardial Damage Following Coronary Microembolization in Dogs. Acta Pathologica Microbiologica et Immunologica Scandinavica - Section A Pathology 1983, 91 (2), 115–124.

(11) Sabbah, H. N.; Stein, P. D.; Kono, T.; Gheorghiade, M.; Levine, T. B.; Jafri, S.; Hawkins, E. T.; Goldstein, S. A Canine Model of Chronic Heart Failure Produced by Multiple Sequential Coronary Microembolizations. American Journal of Physiology - Heart and Circulatory Physiology 1991, 260 (4), H1379–84.

(12) Firoozan, S.; Wei, K.; Linka, A.; Skyba, D.; Goodman, N. C.; Kaul, S. A Canine Model of Chronic Ischemic Cardiomyopathy: Characterization of Regional Flow-Function Relations. American Journal of Physiology - Heart and Circulatory Physiology 1999, 276 (2), H446–H455.

(13) Adamson, P. B.; Vanoli, E. Early Autonomic and Repolarization Abnormalities Contribute to Lethal Arrhythmias in Chronic Ischemic Heart Failure: Characteristics of a Novel Heart Failure Model in Dogs with Postmyocardial Infarction Left Ventricular Dysfunction. Journal of the American College of Cardiology 2001, 37 (6), 1741–1748.

(14) National Research Council. Guide for the Care and Use of Laboratory Animals - Committee for the Update of the Guide for the Care and Use of Laboratory Animals, Institute for Laboratory Animal Research; 2011; Vol. 327.

(15) Basso, C.; Thiene, G.; Della Barbera, M.; Angelini, A.; Kirchengast, M.; Iliceto, S. Endothelin A-Receptor Antagonist Administration Immediately after Experimental Myocardial Infarction with Reperfusion Does Not Affect Scar Healing in Dogs. Cardiovascular Research 2002, 55 (1), 113–121.

(16) Theroux, P.; Ross, J.; Franklin, D.; Covell, J. W.; Bloor, C. M.; Sasayama, S. Regional Myocardial Function and Dimensions Early and Late after Myocardial Infarction in the Unanesthetized Dog. Circulation Research 1977, 40 (2), 158–165.

(17) Lindal, S.; Smiseth, O. A.; MjØS, O. D.; Myklebust, R.; JØRgensen, L. Reversible and Irreversible Changes in the Dog Heart During Acute Left Ventricular Failure Due to Experimental Multifocal Ischemia. Acta Pathologica Microbiologica Scandinavica Series A :Pathology 1986, 94 (3), 177–186.

(18) Corrado, D.; Basso, C.; Judge, D. P. Arrhythmogenic Cardiomyopathy. Circulation Research 2017, 121 (7), 785–802.

(19) Saku, K.; Kakino, T.; Arimura, T.; Sunagawa, G.; Nishikawa, T.; Sakamoto, T.; Kishi, T.; Tsutsui, H.; Sunagawa, K. Left Ventricular Mechanical Unloading by Total Support of Impella in Myocardial Infarction Reduces Infarct Size, Preserves Left Ventricular Function, and Prevents Subsequent Heart Failure in Dogs. Circulation. Heart failure 2018, 11 (5), e004397.

(20) Suzuki, M.; Asano, H.; Tanaka, H.; Usuda, S. Development and Evaluation of a New Canine Myocardial Infarction Model Using a Closed-Chest Injection of Thrombogenic Material. Japanese Circulation Journal 1999, 63 (11), 900–905.

(21) Kim, W. G.; Shin, Y. C.; Hwang, S. W.; Lee, C.; Na, C. Y. Comparison of Myocardial Infarction with Sequential Ligation of the Left Anterior Descending Artery and Its Diagonal Branch in Dogs and Sheep. International Journal of Artificial Organs 2003, 26 (4), 351–357.

(22) He, K. L.; Dickstein, M.; Sabbah, H. N.; Yi, G. H.; Gu, A.; Maurer, M.; Wei, C. M.; Wang, J.; Burkhoff, D. Mechanisms of Heart Failure with Well Preserved Ejection Fraction in Dogs Following Limited Coronary Microembolization. Cardiovascular Research 2004, 64 (1), 72–83.

(23) Arimura, T.; Saku, K.; Kakino, T.; Nishikawa, T.; Tohyama, T.; Sakamoto, T.; Sakamoto, K.; Kishi, T.; Ide, T.; Sunagawa, K. Intravenous Electrical Vagal Nerve Stimulation Prior to Coronary Reperfusion in a Canine Ischemia-Reperfusion Model Markedly Reduces Infarct Size and Prevents Subsequent Heart Failure. International Journal of Cardiology 2017, 227, 704–710.

(24) Gupta, R. C.; Shimoyama, H.; Tanimura, M.; Nair, R.; Lesch, M.; Sabbah, H. N. SR Ca2+-ATPase Activity and Expression in Ventricular Myocardium of Dogs with Heart Failure. American Journal of Physiology - Heart and Circulatory Physiology 1997, 273 (1), H12–18.

(25) Sabbah, H. N.; Shimoyama, H.; Kono, T.; Gupta, R. C.; Sharov, V. G.; Scicli, G.; Levine, T. B.; Goldstein, S. Effects of Long-Term Monotherapy with Enalapril, Metoprolol, and Digoxin on the Progression of Left Ventricular Dysfunction and Dilation in Dogs with Reduced Ejection Fraction. Circulation 1994, 89 (6), 2852–2859.

(26) Morita, H.; Khanal, S.; Rastogi, S.; Suzuki, G.; Imai, M.; Todor, A.; Sharov, V. G.; Goldstein, S.; O’Neill, T. P.; Sabbah, H. N. Selective Matrix Metalloproteinase Inhibition Attenuates Progression of Left Ventricular Dysfunction and Remodeling in Dogs with Chronic Heart Failure. American Journal of Physiology - Heart and Circulatory Physiology 2006, 290 (6), H2522–227.

